# Development of a Complement Hemolysis Assay Using Aldehyde-Modified Human Erythrocytes

**DOI:** 10.64898/2026.06.16.732604

**Authors:** Brian Andrich L. Pollo, Ruby Anne N. King, Fresthel Monica M. Climacosa, Salvador Eugenio C. Caoili

## Abstract

**Background:** Complement-mediated hemolysis assays are essential for assessing immune function and diagnosing complement-related disorders. Conventional human erythrocyte derivatization with 2,4,6-trinitrobenzene sulfonic acid (TNBS) can induce nonspecific hemolysis and optical interference, complicating interpretation. Identifying a more biocompatible electrophile could improve assay specificity and reliability.

**Methods:** A panel of aldehydes was screened for electrophilicity using a nucleophile consumption assay with glycine as a model nucleophile. Glyoxylic acid was selected based on reactivity, solubility, and visual neutrality, then neutralized with sodium bicarbonate to minimize baseline hemolysis. Human erythrocytes were sequentially treated with pancreatin and glyoxylic acid to generate glyoxylic acid–pancreatin–treated erythrocytes (GxPEs). Complement-mediated hemolysis was assessed using normal human serum, heat-inactivated serum, and pathway-specific conditions, with CH50 values calculated for total, alternative, and non-alternative pathways.

**Results:** GxPEs exhibited robust complement-specific hemolysis (maximum 93.56%) with negligible background activity in heat-inactivated serum. CH50 analysis confirmed activation via both alternative (0.9514 μL) and non-alternative (1.963 μL) pathways. Reconstitution experiments with factor B–depleted cryoprecipitate and cryosupernatant fractions demonstrated dependence on small complement components such as C2 and C4.

**Conclusions:** Glyoxylic acid derivatization yields a reproducible, optically quiet, and complement-specific erythrocyte substrate suitable for functional hemolysis assays. This method offers a practical platform for complement diagnostics, research applications, and therapeutic evaluation.

## BACKGROUND

The complement system is a key component of innate immunity, and its activation is a key feature of various physiologic and pathologic processes such as inflammation, pathogen clearance, and immune complex deposition [1–4]. Traditional methods of assessing complement activity, such as animal erythrocyte hemolysis assays or ELISA-based C3/C5 cleavage assays, are complex, expensive, and time-consuming [5–7]. There is a need therefore for simple, rapid, and cost-effective assays to detect complement activation.

Traditional hemolysis assays rely on animal-derived erythrocytes, such as those obtained from sheep or rabbit, as indicator cells [8,9]. However, sourcing, handling, and ethical clearance for animal-derived materials present logistical and regulatory challenges, especially in countries like the Philippines where access to approved laboratory animals, certified suppliers, and accredited animal facilities is limited. These difficulties increase the cost, time, and complexity of assay development and validation, underscoring the need for alternatives.

While human erythrocytes have been proposed as one of said alternatives, modification requires complex reagents, and the underlying mechanism of lysis activation favors calcium-dependent pathways (*i.e*., classical and lectin pathways) [10–14]. One such reagent used to derivatize human erythrocytes is 2,4,6-trinitrobenzenesulfonate (TNBS) [15]. TNBS is classified as a hazardous chemical due to its strong electrophilic nature and potential for causing severe skin, eye, and respiratory irritation. It is also an allergen and can induce immune responses (haptenization). This makes TNBS incompatible with large-scale, routine, or clinical reagent preparation. TNBS is classified as a nitroaromatic compound, raising concerns for environmental safety and waste disposal. Further, regulatory hurdles have complicated the potential use of TNBS for diagnostic or pharmaceutical applications.

There is a need therefore for alternative indicator cells that circumvents animal and TNBS dependency. Such cells would address the reproducibility, sourcing, and ethical limitations of existing methods, particularly in regions with restricted access to animal-derived materials and facilities [16,17]. The present study addresses this gap by proposing a novel platform for complement detection based on human, non-toxic indicator cells.

## METHODS

### Nucleophile Consumption Assay

To identify an alternative electrophile to TNBS for erythrocyte derivatization, a panel of candidate aldehydes was screened. Electrophilicity screening was conducted using a nucleophile consumption assay. Said assay comprises the following steps: (1) Candidate aldehyde and glycine (as model nucleophile) are reacted in a Schiff base (imine) formation reaction; (2) Adding TNBS to back-titrate excess unreacted glycine; and (3) Measuring the resulting yellow-colored product at 450nm.

Specifically, said assay is briefly described as follows: A 50-μL volume of 3mM glycine in 0.9% saline solution (NSS), pH 7.0, was placed in each well of a 96-well flat-bottom plain polystyrene plate. A 50-uL volume of 2-fold serial dilution of candidate aldehydes in NSS was then added, and the resulting solution incubated at 37°C for 1hr. Excess unreacted glycine was then tested by the addition of 50μL 3mM TNBS in NSS. The optical density at 450nm (OD450nm) was measured after incubation at 37°C for 1hr.

### Glyoxylic Acid Neutralization

To rule out the possibility of pH contributing to baseline hemolysis, various solutions of glyoxylic acid were tested in the absence of complement. The following glyoxylic acid solutions were tested: Glyoxylic acid diluted in NSS (FIG. 2A), glyoxylic acid diluted in CBS (FIG. 2B), glyoxylic acid neutralized with 5N NaOH to pH 7, then diluted with NSS (FIG. 2C), and glyoxylic acid neutralized with 1M NaHCO3 to pH 8.5, then diluted with CBS (FIG. 2D).

One part erythrocyte was treated with 4 parts of 2-fold serial dilution of glyoxylic acid solution (starting 540mM), then diluted with the corresponding buffer for a final erythrocyte concentration of 1% (v/v). Hemolysis was measured as change in OD650nm, taken every 5 minutes, starting 2 minutes after addition of glyoxylic acid solution, up to 60 minutes.

### Complement Hemolysis Assay

A complement-mediated hemolysis assay was developed from human erythrocytes chemically derivatized with glyoxylic acid and enzymatically hydrolyzed with pancreatin. Preparatory steps prior to said complement-mediated hemolysis assay include: (1) Erythrocyte isolation, and (2) erythrocyte modification. Erythrocytes were isolated from whole blood by centrifugation at 1500g for 5 mins. Washing was done thrice with NSS prior to subsequent treatment.

For erythrocyte modification, a volume of erythrocytes was incubated with 4 volumes of a 2-fold serial dilution of aldehyde in buffer. The starting concentration was the highest achievable concentration based on solubility. The buffer used was NSS unless otherwise specified. Incubation of the erythrocyte-aldehyde suspension was then conducted in 37°C for 1 hour. Washing was once again performed thrice with NSS prior to use in assays.

Erythrocyte modification involved sequential treatment with pancreatin and glyoxylic acid or TNBS. Pancreatin treatment involved incubating for 4 hours at 37°C a volume of packed erythrocytes with 5 volumes of 5%(w/v) pancreatin in phosphate-buffered saline (pH 7.4). Glyoxylic acid treatment involved incubating for 1 hour at 37°C a volume of packed erythrocytes with 4 volumes of 540mM glyoxylic acid neutralized with 1M NaHCO_3_ and diluted in CBS, as previously optimized. TNBS treatment involved incubating for 1.5 hours at 4°C a volume of packed erythrocytes with 4 volumes of 2mM TNBS in CBS. Washing was done thrice with NSS in between each treatment. The order of treatment was tested as well.

The susceptibility of the modified erythrocytes to complement was tested using the developed complement-mediated hemolysis assay. In brief, a sufficient volume of said modified erythrocytes was diluted with HEPES-buffered saline, pH 7.4, containing 15mM 4-(2-hydroxyethyl)-1-piperazineethanesulfonic acid (HEPES) and 150mM NaCl, supplemented with 0.3mM Ca^2+^ and 2mM Mg^2+^ (HBS++), a buffer capable of supporting complement activation. Said modified erythrocytes were diluted for a final assay concentration of 1%. A 50-μL volume of each suspension was placed in each well, then 50μL of 1:1 normal human sera (NHS)-HBS++ was added. The NHS used in all experiments is a pooled sample representing 20 individual healthy donors. After incubation at 37C for 1 hour, optical density was measured at 650nm. Percent hemolysis was calculated relative to negative and positive controls using HBS++ and 1% Triton X, respectively, instead of NHS-HBS++. Heat-inactivated sera (HIS) was used as control for complement source in place of NHS. HIS was prepared by incubating NHS in 56°C for 30 min.

### Pathway-Specific Complement Hemolysis Assay

NHS was depleted of certain complement components to favor the activation of a particular pathway. For instance, R(B) is NHS depleted of factor B by incubating in 50°C for 20 minutes, thus favoring the activation of non-alternative pathways. To favor the activation of the alternative pathway, Mg-EGTA was used, which is NHS relatively depleted of Ca^2+^ and enriched in Mg^2+^, by addition of 8mM EGTA and 10mM MgCl_2_. EGTA is a calcium-specific chelator. The relative enrichment of Mg^2+^ and depletion of Ca^2+^ ions disfavors the calcium-dependent pathways of complement activation (*i.e.,* the classical and lectin pathways). When Mg-EGTA was used as complement source, HBS-EGTA was the buffer used in place of HBS++. HBS-EGTA is HEPES-buffered saline containing 8mM EGTA and 10mM MgCl_2_. NHS was used to evaluate the effect of activation of the total complement system (*i.e.,* all pathways active), while HIS served as negative control.

Cryoprecipitation was used to fractionate the complement components of fresh frozen plasma (FFP) by relative size. Cryoprecipitate (CryoP) is relatively enriched in larger complement components such as C1q and mannose-binding lectin (MBL). In contrast, cryosupernatant (CryoS) retains other smaller complement components. Cryoprecipitation was performed by thawing 1 unit of FFP overnight (12 hours) at 4 ± 2°C. The precipitate thus obtained was designated as CryoP, while the supernatant was designated as CryoS. Prior to use in assays, CryoP was resuspended in HBS++ at a volume equal to the original source FFP. In the experiment depicted in FIG. 5, CryoP and CryoS were depleted of factor B by heating at 50°C for 20 min prior to use in the assay. Such fractions were used as alternative complement source in place of NHS. **RESULTS**

### Aldehyde Screening

To identify an alternative electrophile to TNBS for erythrocyte derivatization, a panel of candidate aldehydes was screened.

Screening revealed minimal glycine reactivity and electrophilicity of glucose (with sorbitol as comparator), benzaldehyde, and vanillin (FIGS. 1A,C,D). In contrast, glyoxylic acid and *ortho-*phthalaldehyde consumed glycine at a median concentration of 26.62mM (95%CI: 19.22 – 30.34) and 5.13mM (95%CI: 4.72 – 5.28), respectively. To explore the reaction buffer, said nucleophile consumption assay was performed with carbonate-buffered saline (CBS), pH 8.5, containing 120mM NaHCO3 and 40mM NaCl, to favor the deprotonated state of primary amines, thus theoretically improving reactivity towards the aldehydic electrophiles. However, no significant change in reactivity was observed with vanillin and glyoxylic acid, suggesting that the reaction pH is not crucial in the tested aldehyde-glycine reactions.

To explore the effect of prior neutralization, glyoxylic acid was neutralized with either 5N NaOH or 1M NaHCO_3_. The obtained hemolysis curves depicted in FIG.1E suggest neutralization with 1M NaHCO_3_ attenuates nucleophile consumption, while neutralization with 5N NaOH accelerates nucleophile consumption.

Given its ready availability, high solubility, and colorless nature, glyoxylic acid was pursued in subsequent experiments as the leading candidate aldehyde.

### Glyoxylic acid neutralization

Nonspecific hemolysis was observed with higher dilutions of non-neutralized glyoxylic acid, suggesting the need for neutralization prior to use in erythrocyte modification. Neutralization with NaOH led to excess osmolarity as well as precipitation. It also contributed to an increase in baseline turbidity. In contrast, glyoxylic acid neutralized with NaHCO_3_ did not exhibit hemolysis under the conditions tested, hence this was the neutralization method employed in subsequent studies.

### Complement Hemolysis Assay

The time to reach 50% maximal hemolysis (t_50_) for glyoxylic acid-pancreatin-treated erythrocytes (GxPEs) treated with 25% NHS was 7.656 minutes, with a maximum achieved hemolysis of 93.56%. No hemolysis was observed with HIS. This suggests an improvement in assay characteristics over TNBS-pancreatin-treated erythrocytes (TxPEs). While TxPEs treated with NHS achieved 95.12% hemolysis, TxPEs treated with HIS also reached 95.25% hemolysis, suggesting nonspecific hemolysis not mediated by complement, which is a heat-labile system. Nevertheless, the presence of complement appears to hasten hemolysis, with t_50_ for TxPEs treated with NHS being 9.921 minutes (95%CI: 9.510 – 10.240 min), while t_50_ for TxPEs treated with HIS being 14.44 minutes (95% CI: 14.09 – 14.80 min). Order of treatment also appears to be important, as treatment with pancreatin first led to extensive hemolysis in the case of subsequent TNBS treatment, and highly variable hemolysis in the case of subsequent glyoxylic acid treatment.

### Complement Hemolysis Pathways

To probe the specific pathways involved in complement-mediated hemolysis of modified erythrocytes, variations of the complement hemolysis assay were performed.

The CH50 determined for total activation, the non-alternative pathways, and the alternative pathway were 2.447μL (95%CI: 2.148 – 2.772), 1.963 μL (95%CI: 1.763 – 2.188), and 0.9514 μL (95%CI: 0.8966 – 1.009), respectively. This suggests that the assay detects hemolysis mediated by alternative and non-alternative pathways alone and/or in concert. This contrasts with prior complement-mediated hemolysis assays that have a preference for non-alternative pathways.

R(B)CryoP, which is depleted of the smaller complement components, had low hemolytic potency, with a CH_50_ of 7.43μL. Restoration of these components by admixing R(B)CryoP with R(B)CryoS reconstituted hemolytic potency. R(B)CryoS has a CH50 of 0.2827μL (95%CI: 0.2482 – 0.3138), while R(B)CryoP+ R(B)CryoS at a 1:1 ratio has a CH50 of 0.4307μL (95%CI: 0.3298 – 0.5784). Overall, these findings are consistent with assay dependence on the smaller complement components (*i.e.*, C2, and C4) involved in the non-alternative pathways.

## DISCUSSION

This study introduces glyoxylic acid as a viable alternative to TNBS for erythrocyte derivatization in complement hemolysis assays. Glyoxylic acid, when neutralized with sodium bicarbonate, provided a non-hemolytic, colorless, and reactive aldehyde suitable for modifying erythrocytes without baseline turbidity or osmotic disruption. The resulting GxPEs showed strong complement-mediated hemolysis with low background activity in HIS. Furthermore, GxPEs were sensitive to both alternative and non-alternative complement pathways, as shown by CH50 analysis, including experiments using reconstituted cryoprecipitate-depleted sera.

Human erythrocyte derivatization is a proposed alternative to the traditional strategies for probing complement activity using nonhuman animal erythrocytes [18,19]. One such derivatization strategy involves TNBS [20]. However, TNBS’s strong chromophoric properties and potential to provoke non-complement-mediated hemolysis complicate interpretation. Here, glyoxylic acid offered distinct advantages. While aldehydes such as vanillin and benzaldehyde showed minimal electrophilicity in our glycine assay, consistent with prior reactivity studies, the high solubility and reactivity at physiological pH of glyoxylic acid made it uniquely suitable for erythrocyte derivatization. We speculate that the mechanism of derivatization includes at least the classical reaction of glyoxylic acid with tryptophan known as the Hopkins-Cole reaction [21].

The improved specificity of GxPE hemolysis to complement was confirmed by the absence of lysis in HIS-treated cells, contrasting with TxPEs, which showed residual hemolysis suggestive of chemical instability or membrane fragility independent of complement. The faster t50 for GxPEs compared to TxPEs also suggests improved surface presentation of complement-activating epitopes, potentially due to more uniform modification.

The data implicate both classical/lectin (non-alternative) and alternative complement pathways in hemolysis of GxPEs. This distinguishes the current assay from the classical CH50 assay, which is largely insensitive to factor B depletion [20]. The restoration of hemolytic activity by admixing R(B)CryoS with R(B)CryoP indicates dependency on small complement components (e.g., C2, C4), further affirming the activation of the classical and/or lectin pathways. The elevated hemolytic capacity of GxPEs relative to earlier models may also reflect pancreatin-induced surface remodeling, which exposes neoepitopes for complement activation, a process reported in limited prior work [22,23] but not well explored with non-TNBS reagents.

One limitation was the lack of direct structural characterization of modified erythrocytes. Surface proteomics or glycomics may confirm the nature of modifications introduced by glyoxylic acid. Furthermore, while the glycine consumption assay provides a convenient surrogate for aldehyde reactivity, it may not fully predict erythrocyte surface reactivity due to the complexity of membrane proteins and their steric environment. Lastly, while NaOH neutralization enhanced glycine reactivity, its associated hemolysis and precipitation precluded practical use, suggesting that there is a tradeoff between chemical reactivity and biocompatibility.

The shift away from TNBS-based assays offers a more chemically benign, optically silent alternative. Previous derivatization strategies often required reduction steps (e.g., using NaBH4) to stabilize Schiff bases [24], whereas glyoxylic acid may form stable adducts without reduction under our conditions. Unlike OPA, which is fluorescent and forms isoindole products in presence of thiols [25–27], glyoxylic acid’s behavior in our assay system provides a neutral baseline and avoids signal interference. Furthermore, unlike dextran or immunoglobulin-coated erythrocytes, which bias toward classical pathway activation [28–30], GxPEs show more balanced pathway sensitivity.

This platform is proof-of-concept for a more versatile complement hemolysis assay that could be tuned toward specific pathway interrogation. Future work could explore detailed surface chemistry as well as assess assay performance across donor variability and complement deficiencies. The simplicity and reproducibility of glyoxylic acid derivatization also make it an attractive candidate for diagnostic or screening assays in complement research, including complement deficiencies, autoimmune disorders, or the evaluation of complement-targeting therapeutics.

## CONCLUSION

Glyoxylic acid, neutralized with sodium bicarbonate, is a practical and chemically benign alternative to TNBS for human erythrocyte derivatization in complement hemolysis assays. The resulting GxPEs exhibit minimal background lysis, robust complement specificity, and balanced pathway sensitivity, providing a reproducible platform for functional complement testing. This simple derivatization method enables broader assay versatility and potential diagnostic and therapeutic applications.

## ACKNOWLEDGEMENTS

The authors would like to express their appreciation to the Department of Science and Technology Philippine Council for Health Research and Development (DOST-PCHRD) for providing support through scholarships and dissertation grants. The Section of Blood Banking and Transfusion Medicine, Department of Laboratories, University of the Philippines - Philippine General Hospital (UP-PGH) provided blood samples for testing.

## DATA AVAILABILITY STATEMENT

Raw data are available in the Harvard Dataverse (https://doi.org/10.7910/DVN/EALBPP). More information on the methods are available in protocols io (https://dx.doi.org/10.17504/protocols.io.q26g77ye3gwz/v1).

---

**Figure 1.**
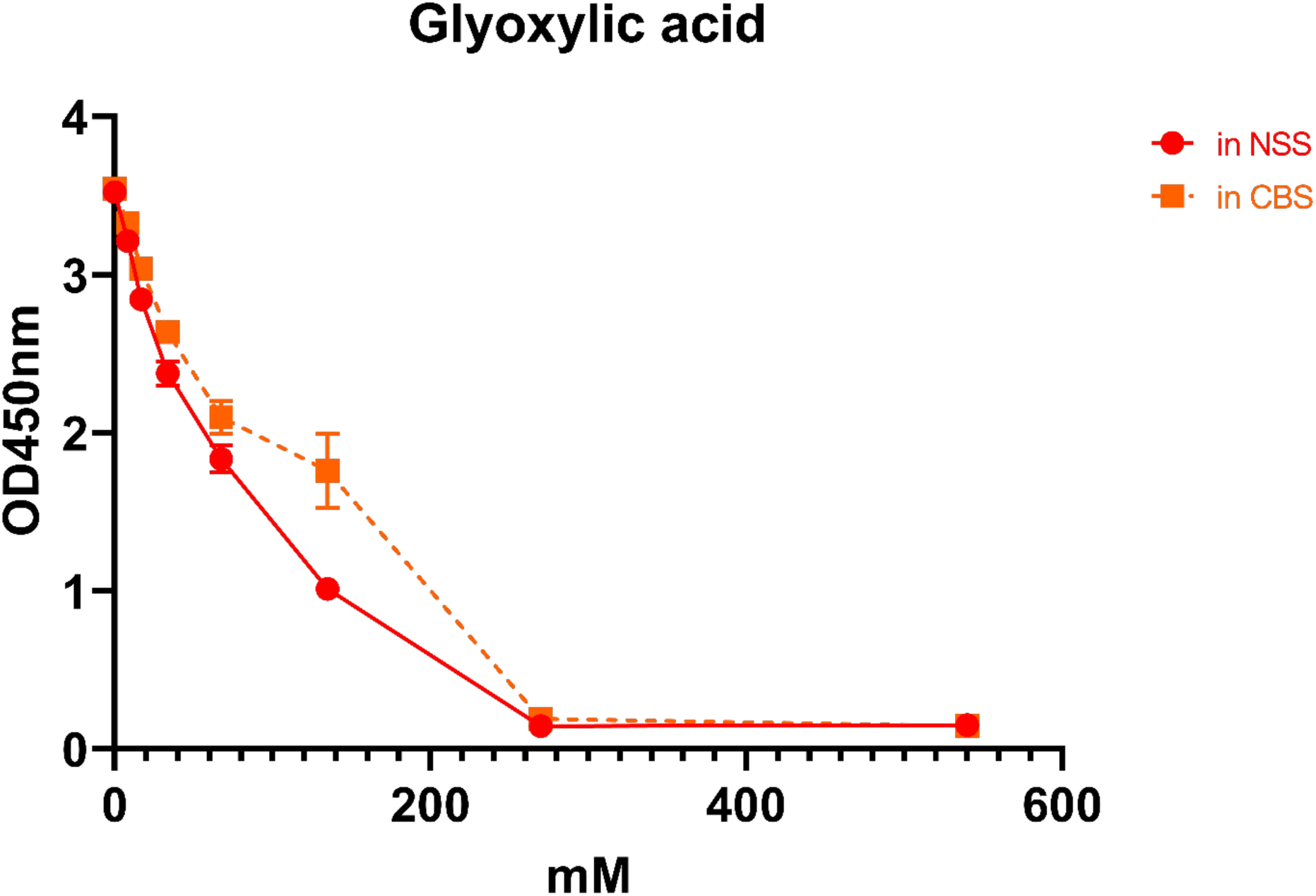
Electrophilicity of glyoxylic acid was demonstrated using a nucleophile consumption assay. Said assay comprises the steps of: contacting said candidate aldehyde with a nucleophile (*i.e.*, glycine), then back-titrating remaining excess unreacted nucleophile with TNBS, with the resulting trinitrophenylated glycine measured at OD450nm; data represent mean ± SD (*n* = 3); OD450nm: optical density at 50nm; NSS: 0.9% saline solution (pH 7); CBS: carbonate-buffered saline solution (pH 8.5).

**Figure 2.**
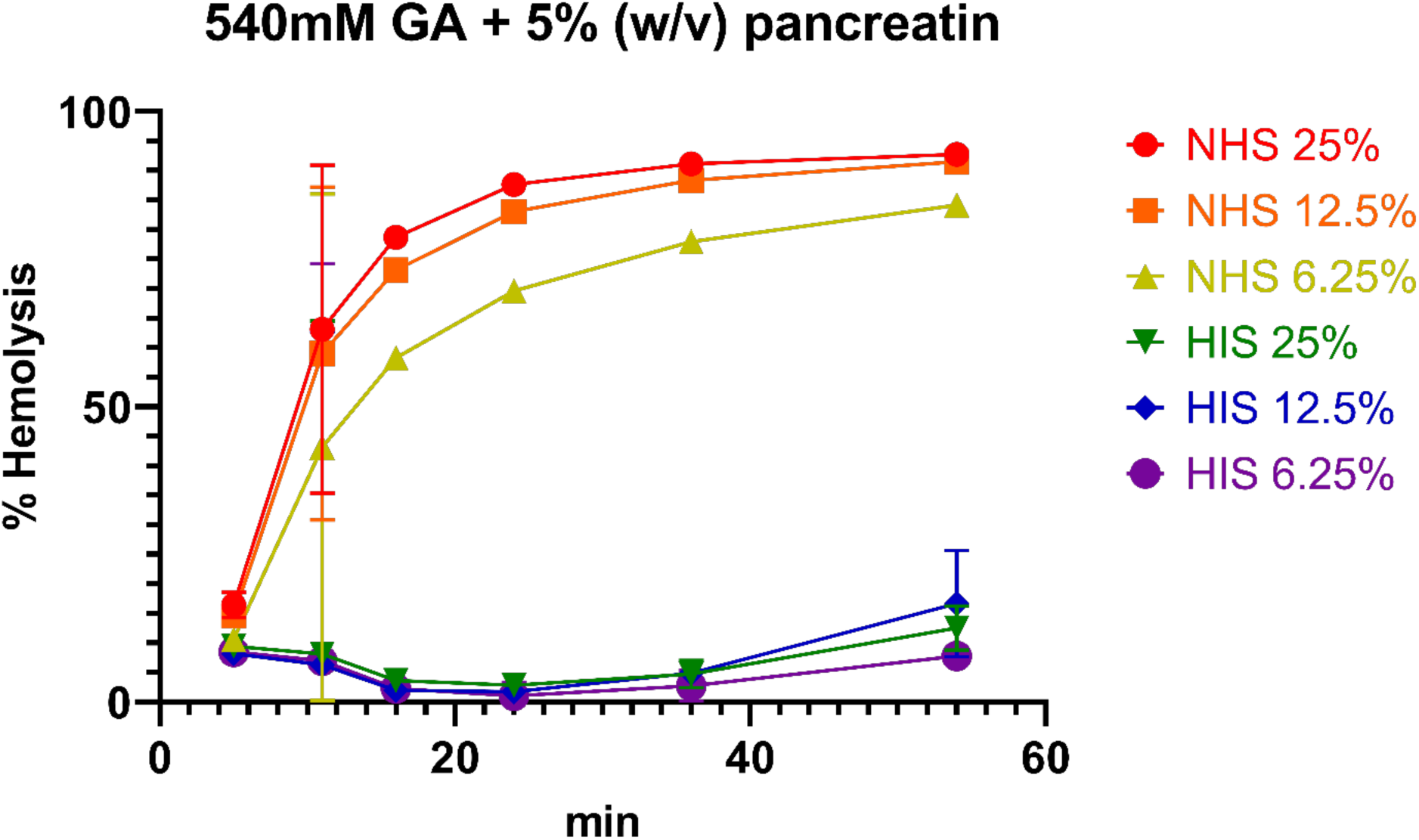
Sequential treatment of human erythrocytes with glyoxylic acid and pancreatin renders human erythrocytes susceptible to complement. Complement-mediated hemolysis assays were performed to determine the effect of treatment with glyoxylic acid compared to TNBS, before or after treatment with pancreatin, on the susceptibility of erythrocytes to NHS as complement source and HIS as control; erythrocytes were treated for 1 hour in 37°C with 2-fold serial dilution of various forms of glyoxylic acid solution starting from 540 mM, then hemolysis measured as a decrease of OD650nm relative to control; hemolysis was measured on a logarithmic timescale starting at 5 minutes after addition of glyoxylic acid solution (11, 16, 24, 36, 54 min); data represent mean ± SD (*n* = 3); NHS: pooled normal human sera (*n* = 20); HIS: NHS heat-inactivated by incubating in 56°C for 30 minutes; TNBS: 2,4,6-trinitrobenzenesulfonate; GA: glyoxylic acid.

**Figure 3.**
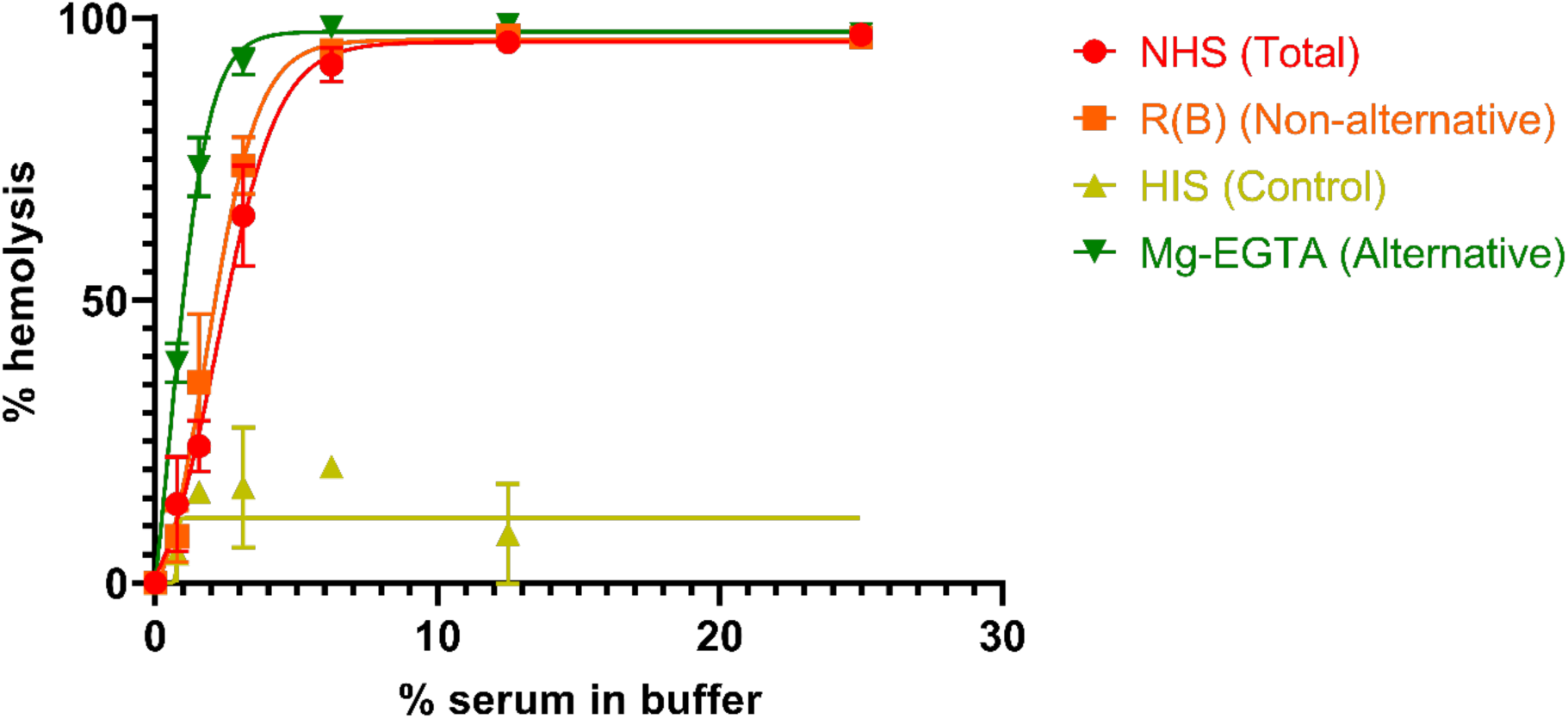
Complement-mediated hemolysis assays demonstrate that erythrocytes sequentially treated with glyoxylic acid and pancreatin are susceptible via both alternative and non-alternative pathways. the complement sources are NHS for total complement activation, R(B) for non-alternative pathways activation, Mg-EGTA for alternative pathway activation, and HIS as control; data represent mean ± SD (*n* = 3); regression line corresponds to 4-parameter logistic regression; NHS: pooled normal human sera (*n* = 20); R(B): NHS depleted of factor B by incubating in 50°C for 20 minutes; Mg-EGTA: NHS relatively depleted of Ca^2+^ and enriched in Mg^2+^, by addition of 8mM EGTA and 10mM MgCl_2_; HIS: NHS heat-inactivated by incubating in 56°C for 30 minutes; EGTA: ethylene glycol-bis(2-aminoethylether)-*N,N,N′,N′*-tetraacetic acid.

**Figure 4.**
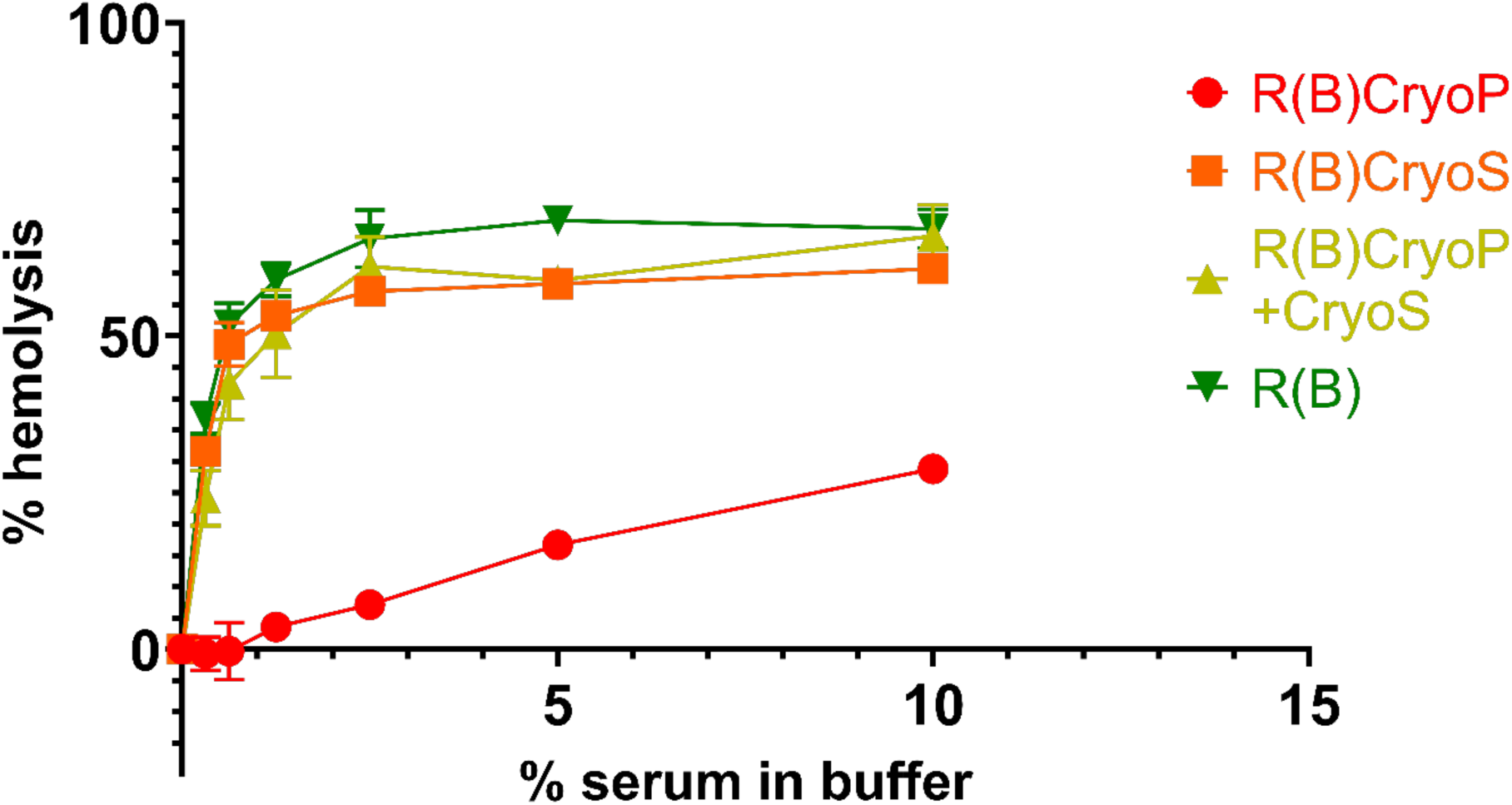
The addition of removed complement factors from plasma restores hemolysis; R(B) was utilized to favor the non-alternative pathways, while cryoprecipitation was used to fractionate complement components into larger (CryoP) and smaller components (in the CryoS); data represent mean ± SD (*n* = 3); R(B): NHS depleted of factor B by incubating in 50°C for 20 minutes; CryoP: the precipitate obtained upon thawing FFP at 4 ± 2°C; CryoS: the supernatant obtained upon thawing FFP at 4 ± 2°C.

**Supplementary Figure 1.**
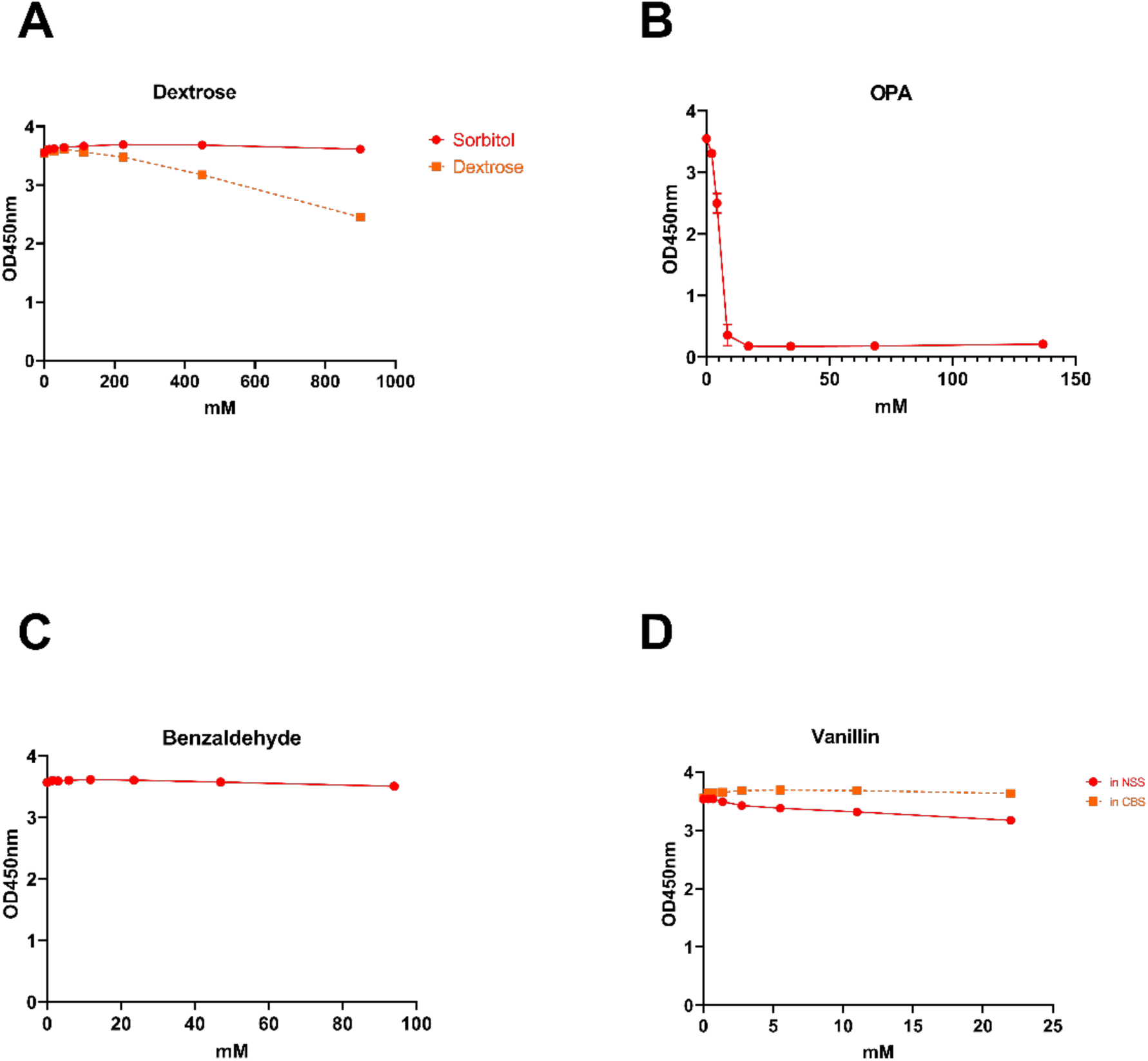
A panel of candidate aldehydes were screened as TNBS substitutes; Said candidate aldehydes were screened for electrophilicity using a nucleophile consumption assay; Said assay comprises the steps of: contacting said candidate aldehyde with a nucleophile (*i.e.*, glycine), then back-titrating remaining excess unreacted nucleophile with TNBS, with the resulting trinitrophenylated glycine measured at OD450nm; data represent mean ± SD (*n* = 3); OD450nm: optical density at 450nm; OPA: *ortho-*phthalaldehyde; NSS: 0.9% saline solution (pH 7); CBS: carbonate-buffered saline solution (pH 8.5); MBS: sodium metabisulfite.

**Supplementary Figure 2.**
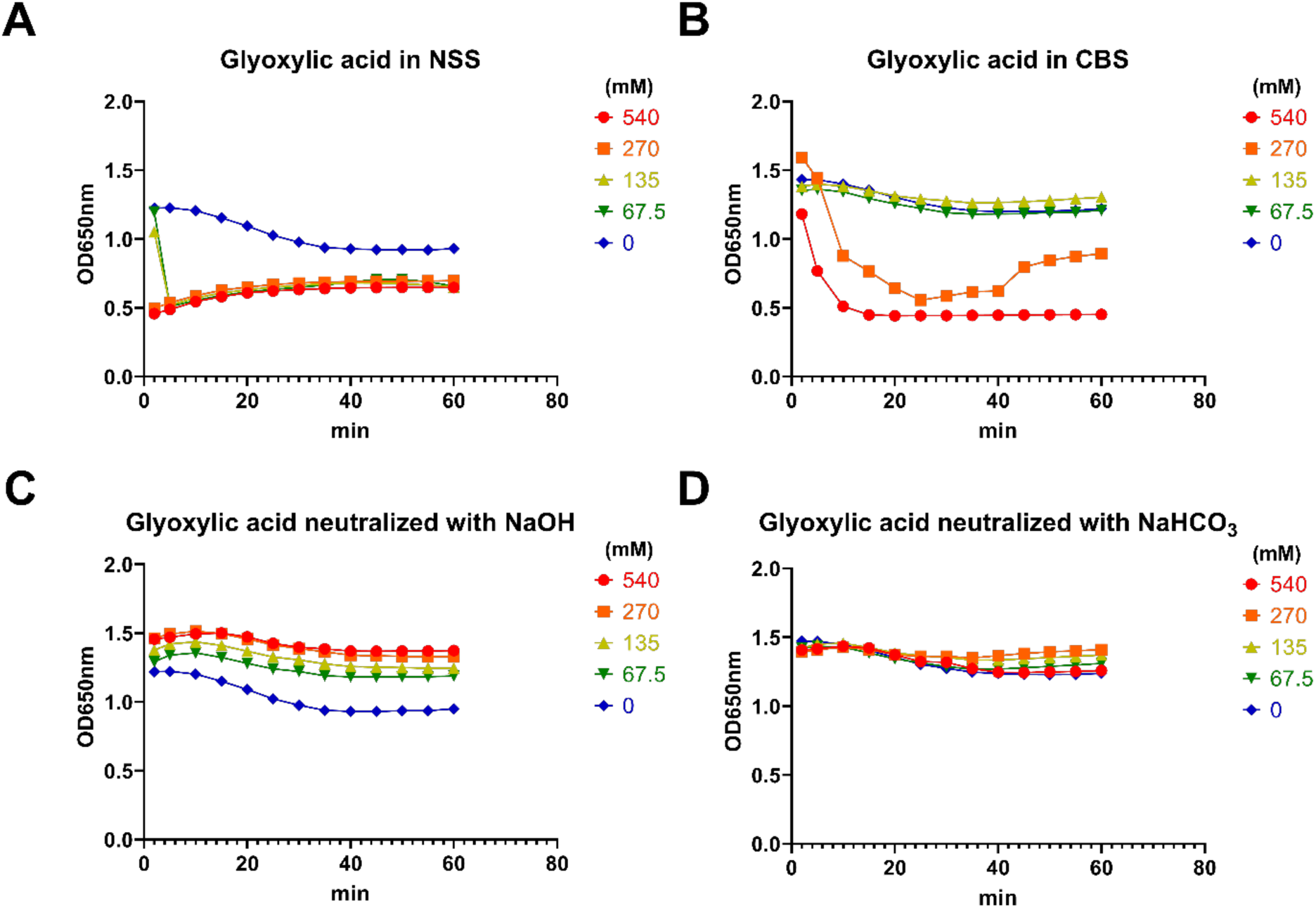
Kinetic hemolysis assays were performed to determine the effect of neutralization on the baseline hemolytic activity of glyoxylic acid, using NaOH and NaHCO_3_ as neutralizing agents; erythrocytes were treated for 1 hour in 37°C with 2-fold serial dilution of various forms of glyoxylic acid solution starting from 4mM, then hemolysis measured as a decrease of optical density at 650nm; hemolysis was measured every 5 minutes starting at 2 minutes after addition of glyoxylic acid solution; *n* = 1; OPA: *ortho-*phthalaldehyde; NSS: 0.9% saline solution (pH 7); CBS: carbonate-buffered saline solution (pH 8.5).

**Supplementary Figure 3.**
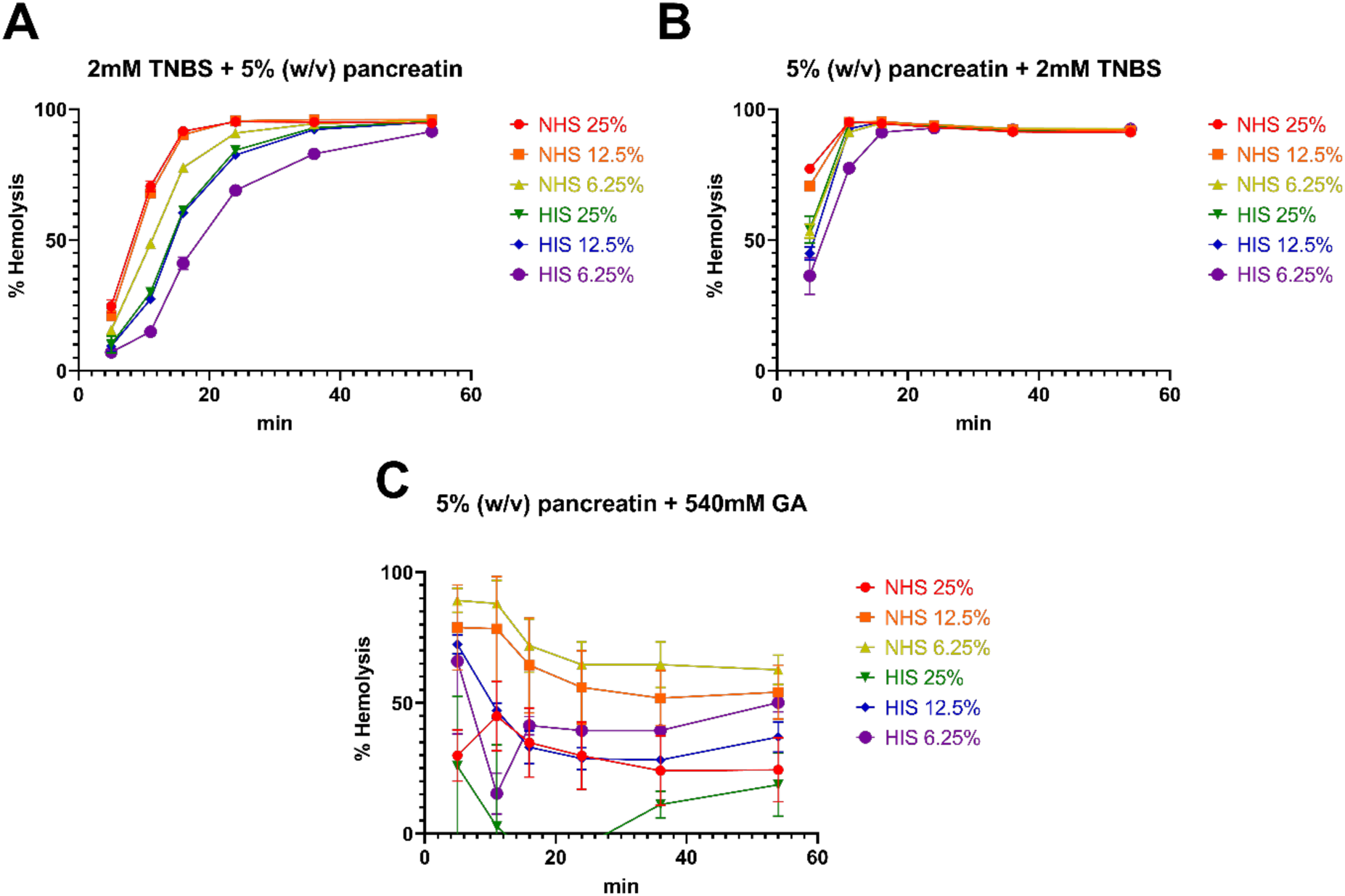
Complement-mediated hemolysis assays were performed to determine the effect of treatment with glyoxylic acid compared to TNBS, before or after treatment with pancreatin, on the susceptibility of erythrocytes to NHS as complement source and HIS as control; erythrocytes were treated for 1 hour in 37°C with 2-fold serial dilution of various forms of glyoxylic acid solution starting from 540 mM, then hemolysis measured as a decrease of OD650nm relative to control; hemolysis was measured on a logarithmic timescale starting at 5 minutes after addition of glyoxylic acid solution (11, 16, 24, 36, 54 min); data represent mean ± SD (*n* = 3); NHS: pooled normal human sera (*n* = 20); HIS: NHS heat-inactivated by incubating in 56°C for 30 minutes; TNBS: 2,4,6-trinitrobenzenesulfonate; GA: glyoxylic acid.

